# Age-related differences in spatial memory occur alongside reduced visual fMRI BOLD but preserved viewpoint-specific scene representations

**DOI:** 10.64898/2026.03.23.713765

**Authors:** Sabina Srokova, Carol A. Barnes, Arne D. Ekstrom

**Affiliations:** Psychology Department, University of Arizona, Tucson, AZ; Evelyn F. McKnight Brain Institute, University of Arizona, Tucson, AZ

## Abstract

Current evidence suggests that older adults perform worse at tasks involving spatial memory and navigation, yet the underlying reasons remain unclear. Here, we tested the hypothesis that age-related declines in spatial memory stem from difficulties in recognizing spatial environments from rotated perspectives. Young and older adults underwent fMRI as they encoded virtual scenes which were later viewed either from the same or rotated perspective. Older adults were worse at identifying changes in these scenes, although the age effect was equally robust across perspective conditions. Neural specificity of scene representations was examined with the phenomenon of fMRI repetition adaptation. We predicted that young adults would show significant fMRI adaptation to the same but not rotated perspective, indicative of intact viewpoint specificity, while older adults show would adaptation effects to both. While analyses of raw fMRI BOLD produced results consistent with these predictions, follow-up analyses revealed a general attenuation of activity in older adults across both perspective conditions. Additionally, although older adults showed both lower fMRI BOLD and worse spatial memory, lower trial-wise BOLD was associated with better performance independent of age. This suggests that the variance associated with fMRI adaptation is reflective of two independent sources of variance: age and cognition. Our results suggest that age differences in spatial memory may manifest due to cognitive and neural factors that are shared across same and rotated perspectives, and thus they cannot be explained by a selective deficit in allocentric (viewpoint-independent) processing.

**Significance Statement:** Increasing age is often associated with reduced spatial memory and navigation. Prior research suggests that age differences in spatial memory could be exacerbated by changes in perspective, possibly due to increased difficulties in the ability to construct allocentric (viewpoint-independent) representations from previously encoded egocentric perspectives. Here, we demonstrate that older adults are equally disadvantaged when recognizing layouts across same and rotated perspectives. FMRI analyses indicate that older age is associated with reduced fMRI BOLD in higher-level visual cortex across both perspective conditions, as opposed to altered specificity of perspective coding. Consequently, the present study challenges the notion that aging is associated with a selective decline in allocentric spatial memory and instead supports a more general age-related difficulty with scene processing.

## Introduction

Compared to young adults, older adults typically perform worse on tasks of spatial memory and navigation (for reviews, see Moffat, 2009; Lester et al., 2017; Coughlan et al., 2018; Hill & Ekstrom, 2025). Although age differences in spatial memory likely reflect multiple contributing factors, evidence suggests that one factor may reflect alterations in the ability to construct viewpoint-independent (allocentric) representations from previously encoded egocentric perspectives (e.g., Bohbot et al., 2012; Lithfous et al., 2014; Castegnaro et al., 2025). In a laboratory setting, individuals’ ability to derive novel viewpoints is typically probed using tasks in which participants encode a visual spatial layout, with their memory for the scene then tested from a rotated perspective. Older adults typically perform worse at these tasks (e.g., King et al., 2002; Montefinese et al., 2015; Muffato et al., 2019; Hilton et al, 2020; Segen et al., 2021), with past work attributing such age differences to alterations in hippocampal-dependent computations. This view is motivated by the putative role of the hippocampus in relational and spatial binding (e.g., King et al., 2002; Hartley et al, 2007; but see Bohbot et al., 1998; McAvan et al., 2022), and the evidence that hippocampal gray matter loss may be accelerated in aging relative to other cortical regions (e.g. Raz et al. 2005, Vidal-Piñeiro et al., 2025). That said, claims attributing age differences in spatial memory to the hippocampus rarely account for the fact that hippocampal computations do not operate in isolation and that they are, in fact, shaped by inputs they receive from cortical regions earlier in the visual hierarchy.

The specificity of neural activity arising in higher-level visual cortex is reduced, or ‘*dedifferentiated*’, in older age, a phenomenon that has been shown to be most robust in the scene-selective visual cortex (for reviews, see Koen & Rugg, 2019; Koen et al., 2020). Neural dedifferentiation is functionally significant to increasing age as it is associated with worse memory for visual stimuli (e.g., Koen et al., 2019, Srokova et al., 2020, 2024, 2025), and with higher levels of cortical tau, a hallmark of Alzheimer’s disease (Maass et al., 2019, Sheng et al., 2025). FMRI adaptation (or suppression), which takes the form of a reduction in the neural response elicited by a repetition of a stimulus, can be used to characterize neural specificity as it reflects reactivation of overlapping neural representations (Grill-Spector et al. 2006, Barron et al., 2016). In the scene perception literature, viewpoint-specific coding is reflective of adaptation effects to scenes of the same viewpoint, but not to rotated perspectives of the same scene (Epstein et al., 2003, 2007, 2008). In the aging literature, fMRI adaptation has been used to demonstrate reduced neural specificity for face and object stimuli, whereby young adults typically show adaptation for exact repetitions only, while older adults often show adaptation for both exact stimulus repeats as well as visually similar stimuli (Goh et al., 2010, Yassa et al., 2011, Reagh et al., 2018). It is possible that age differences in scene processing reflect a shift toward noisier or gist-based viewpoint-invariant coding, accompanied by a degradation of viewpoint-specific coding.

In the present study, young and older adults underwent fMRI as they completed a spatial memory task involving recognizing environments from the same and rotated perspectives. We expected that older age would be associated with reduced ability to recognize spatial layouts, particularly from rotated perspectives. Next, we predicted that while both age groups would exhibit fMRI adaptation to same perspectives, older adults would show elevated adaptation effects to rotated scenes, suggesting that scene representations become less specific with increasing age. Lastly, we expected that attenuated responses to the same perspectives should be predictive of better spatial performance. However, this prediction is less straightforward for rotated perspectives, given that higher adaptation to rotated perspectives, reflective of a shift towards viewpoint invariance, can be both beneficial and detrimental depending on task requirements.

## Materials and Methods

### Participants

Twenty-five young (range = 18 – 28 years, mean (SD) = 22.16 (2.27) years, 13 female) and 27 older adult participants (range = 62 – 78 years, mean (SD) = 69.07 (4.78) years, 17 female) contributed data to the study. One older adult was excluded from analyses as they voluntarily withdrew from the study. Participants were recruited from the University of Arizona and the surrounding areas of Tucson and the Pima county. All participants were compensated for their time at a rate of $20/hour. Participants were right-handed and had normal or corrected-to-normal vision. To ascertain that potential age differences were reflective of normative rather than pathological aging, older adults were included if they had no history of neurological disease, substance abuse, brain injury, diabetes or current/recent use of prescription medication affecting the central nervous system. Additionally, neuropsychological testing was completed to further minimize the likelihood of including participants with mild cognitive impairment or early dementia. The test battery included multiple tests and scores in each of four broad cognitive domains: memory [California Verbal Learning Test-II (CVLT) Long Delay Free Recall (Delis et al., 2000), Rey-Osterrieth Complex Figure Test Long Delay Free Recall, (Rey, 1941)], executive function [Trails A and B, (Reitan & Wolfson, 1985)], language [FAS fluency (Spreen & Benton, 1977), category fluency (Benton, 1968), Boston Naming Test (BNT); Goodglass et al., 2001)], visuo-spatial abilities [Wechsler Adult Intelligence Scale 4th edition (WAIS-IV, Wechsler, 2009), Rey-Osterrieth Complex Figure Test copy score (Rey, 1941)], and verbal intelligence [WAIS-IV Vocabulary and Similarities, (Wechsler, 2009)]. Potential participants were excluded from entry into the study if they scored < 1 SDs below age-appropriate norms on two tests from one domain, or < 1 SDs below age norms on any three tests across all domains (Bondi et al. 2014). All participants provided written informed consent before participation, in accordance with the requirements of the Institutional Review Board of the University of Arizona.

### Experimental Materials and Procedure

Participants completed a spatial working memory task inside an MRI scanner. The task was presented using PsychoPy v2023.2.3 (Pierce et al., 2019), projected onto a display (1920 x 1080 pix. resolution, 41 cm width), placed at the rear end of the scanner bore, and viewed through a mirror mounted on the scanner head coil. The experimental stimuli consisted of 120 image renderings of virtual rooms containing five objects placed in random locations, sized to fit a frame subtending 1067 x 800 pixels. Half of the images were also rendered from a rotated perspective (50-degrees from the center of the room) either to the left or right of the original perspective. To ascertain that each room was perceived as unique, and to reduce across-trial inference, every room was rendered with a unique combination of wall, ceiling, and floor textures. Additionally, the objects within each room were exclusive to that room and did not appear elsewhere in the experiment. Texture and object assets were obtained from the Unity asset store (assetstore.unity.com). Ten additional rooms were generated to be used as practice stimuli prior to the in-scanner task. The stimulus pool described above was used to generate stimulus lists unique to yoked pairs of young and older adults. For all lists, the stimuli were pseudorandomized such that participants did not view more than three consecutive trials of the same condition (perspective or layout, see below). Each list was divided into five blocks (scanner runs) such that each block contained equal number of trials from each condition.

A schematic of the task, including examples of the virtual rooms, are illustrated in **Figure 1**. Participants underwent extensive training and guided practice prior to entering the scanner, which was repeated until the participants showed no confusion and felt comfortable with the task. Following practice, participants were fitted with MR-safe glasses, if necessary, and then went on to complete the experimental task across five scanner runs of 9 minutes and 41 seconds each. Each trial began with a red fixation cross presented for 500 ms, followed by the first presentation of the virtual room for 5000 ms, a pixelated mask for 500 ms, a blue fixation cross for 6500 ms, and then a second presentation of the virtual room for 5000 ms. The second presentation of the room was viewed either from the same perspective, or from a rotated (50-degrees) perspective relative to the first presentation. Additionally, in half of the trials of each perspective condition, one of the five items presented in the room was rendered in a new location. Participants had a total of 5 s to indicate whether the spatial layout was the same (i.e., the objects are in the same locations as before), or different (i.e., one of the objects changed locations). Responses corresponding with the same or different layouts were mapped onto the index and middle fingers of the right hand, which were counterbalanced across participants. The trial ended with a 1000 ms fixation cross, followed by a 3000 – 7000 ms odd-even task. During the odd-even task, participants were shown a double-digit number at the rate of 1000 ms, and their task was to press any button if the presented number was even.

**Figure 1.**
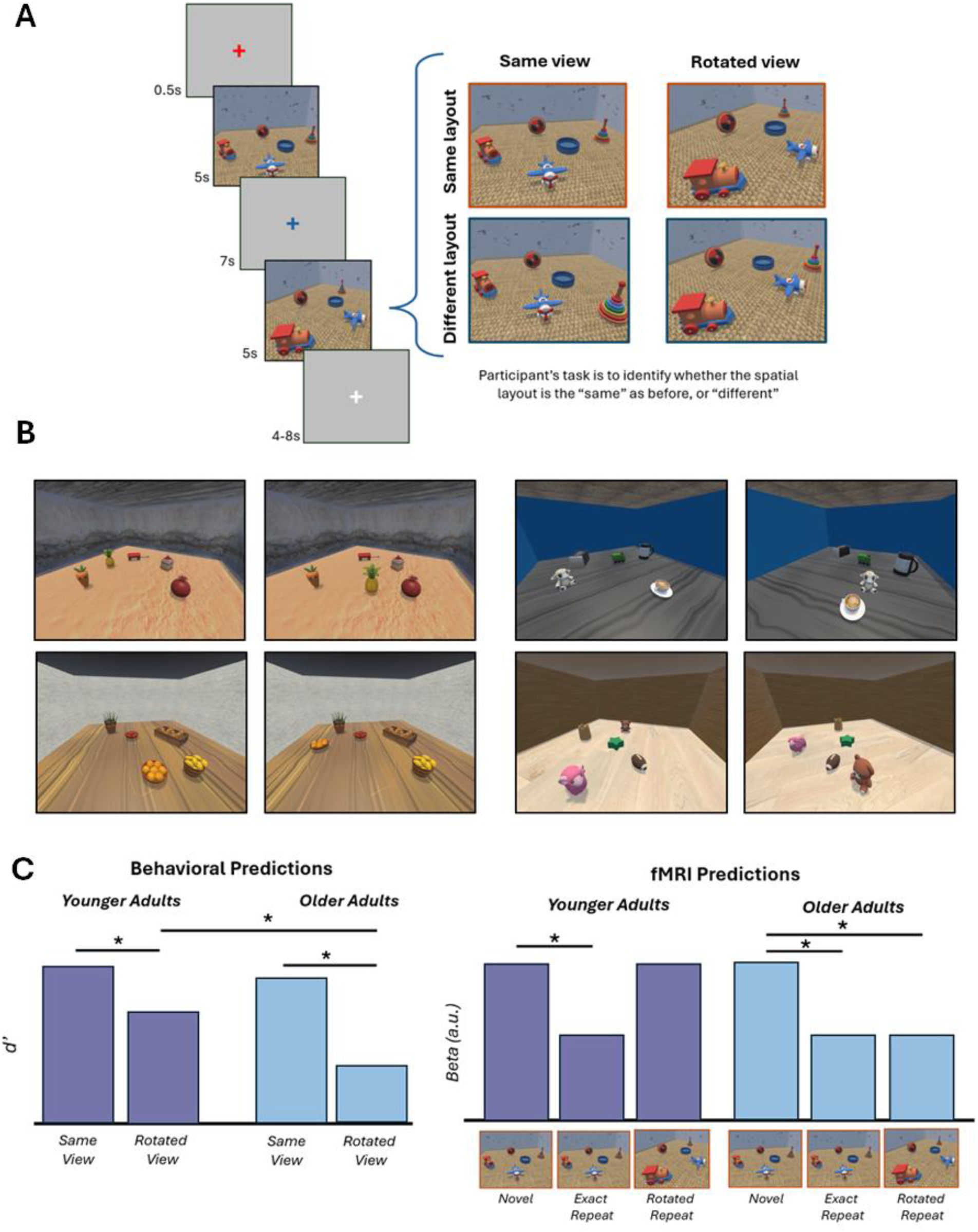
**(A)** Schematic of the experimental paradigm. Participants viewed trial-unique virtual rooms, and after a brief delay, the rooms were presented again either from the same or rotated perspective. On half of the trials, the layout of the objects would change by moving one of the objects to a new location. The participants’ task was to determine if the objects were in the same locations, or to respond ‘different’ if the layout had changed. **(B)** Four example trials which illustrate instances where one of the objects is moved to a new location. Left side = same perspective trials; Right side = rotated perspective. **(C)** Pre-experimental hypotheses. For the behavioral analyses, we predicted that both age groups will perform worse during rotated trials, and that this effect would be disproportionately greater in older adults. For fMRI predictions, we expected that younger adults would exhibit adaptation effects for exact repeats (same perspective, same layout), but not for rotated repeats (rotated perspective, same layout). In contrast, aligning with the hypothesis that higher-level visual regions become less viewpoint-specific with increasing age, older adults were predicted to exhibit adaptation effects for both exact and rotated repeats.

### Behavioral Performance

Participants’ ability to distinguish between trials of same and different spatial layouts was estimated using sensitivity index (d’), calculated as *z(Hit rate) – z(False alarm rate)*, in which hit rate equals the proportion of different layout trials receiving a correct ‘different’ response, and false alarm rate equals proportion of same layout trials receiving an incorrect ‘different’ response. Log-linear correction was applied to both hit and false alarm rates in cases in which either measure equaled 1 or 0, respectively. Response bias/criterion (c), reflecting the amount of evidence required to endorse a spatial layout as different was calculated as: - *0.5 * (z(Hit rate) + z(False alarm rate))*. Estimates of c > 0 reflect a conservative criterion (more frequent ‘same’ responses) and c < 0 reflects a liberal criterion (more frequent ‘different’ responses).

### MRI Data Acquisition and Preprocessing

Functional and structural MRI data were acquired at the University of Arizona using a Siemens Skyra 3T scanner, equipped with a 32-channel head coil. Whole-brain anatomical scan was acquired using a T1-weighted 3D MPRAGE sequence (FOV = 256 × 256 mm; voxel size = 1 × 1 × 1 mm; 160 slices; sagittal acquisition). Functional data were acquired with a T2*-weighted BOLD echoplanar imaging with a multiband factor of 3 (flip angle = 70° FOV = 220 × 220 mm; voxel size = 2 × 2 × 2 mm; TR = 1.52 ms; TE = 30 ms; 66 slices). A dual-echo fieldmap was acquired at 4.92 ms and 7.38 ms immediately after the conclusion of the final task run, resulting in two magnitude images and one pre-subtracted phase image.

The MRI data were preprocessed using Statistical Parametric Mapping (SPM12, Wellcome Department of Cognitive Neurology) and custom MATLAB code (MathWorks). The FieldMap toolbox in SPM was used to calculate voxel displacement maps, which were later applied in SPM’s realign and unwarp procedure. The images were spatially realigned in a two-step procedure: realignment to the first volume of the first scanner run, followed by realignment to the mean EPI image calculated across all scanner runs. Voxel displacement maps were used to perform a dynamic correction of the deformation field. Realigned and unwarped images were manually reoriented along the anterior and posterior commissures, followed by a spatial normalization to an age-unbiased sample-specific EPI template which weighed the contribution of both age groups equally. The normalized data were smoothed with a 5 mm FWHM Gaussian kernel.

### Functional Localizer and Region-of-interest Definition

Following the completion of the in-scanner experimental task, participants completed a functional localizer task for the purpose of delineating functionally defined regions-of-interest (ROIs) independently from the to-be-analyzed data. Participants completed 3 localizer scanner runs, each lasting 4 minutes and 38 seconds. Each scanner run contained 18 randomly interleaved blocks of images of scenes, faces, and objects (6 blocks per image category in each run). A single block contained four exemplars of a given image category, each presented for 2 s and temporally separated by a 1 s fixation cross (extended to 3 s fixation following the last stimulus image of the block). Pleasantness of each image was rated on a 3-point scale to ascertain that participants paid attention to the task. The localizer data was analyzed with a two-stage GLM approach. At the first stage, data for each participant were modeled by convolving a canonical hemodynamic response function (HRF) with a 11 s boxcar onsetting with the first stimulus of a block until the offset of the last image of the block. Regressors were modeled separately for the three image categories. At the second stage, the data were entered into a group-level 2 (age group) x 3 (image category) mixed effects ANOVA.

Participants viewed spatial environments containing object stimuli, and therefore, we conjectured that the task described above engaged visually responsive cortical areas, i.e., visual scene-selective and object-selective cortical regions. We used the data from the functional localizer to define 3 scene-selective regions (scene > object + face contrast): parahippocampal place area (PPA), occipital place area (OPA), retrosplenial complex (RSC), and one object-selective region (object > scene + face contrast): lateral occipital complex (LOC). The contrasts were height thresholded at voxel-wise FWE-corrected p < 0.001 and inclusively masked with anatomical labels provided by the Neuromorphometrics atlas in SPM12. PPA was delimited by the fusiform and parahippocampal gyri, the OPA was masked with the inferior and middle occipital gyri, and the LOC was masked by the inferior occipital, middle occipital, and inferior temporal gyri. RSC is typically not well captured by anatomical labels, and as such, the region was not anatomically constrained and was instead defined by manually separating the cluster from other ROIs. The final *a priori* ROIs are illustrated in **Figure 2**.

**Figure 2.**
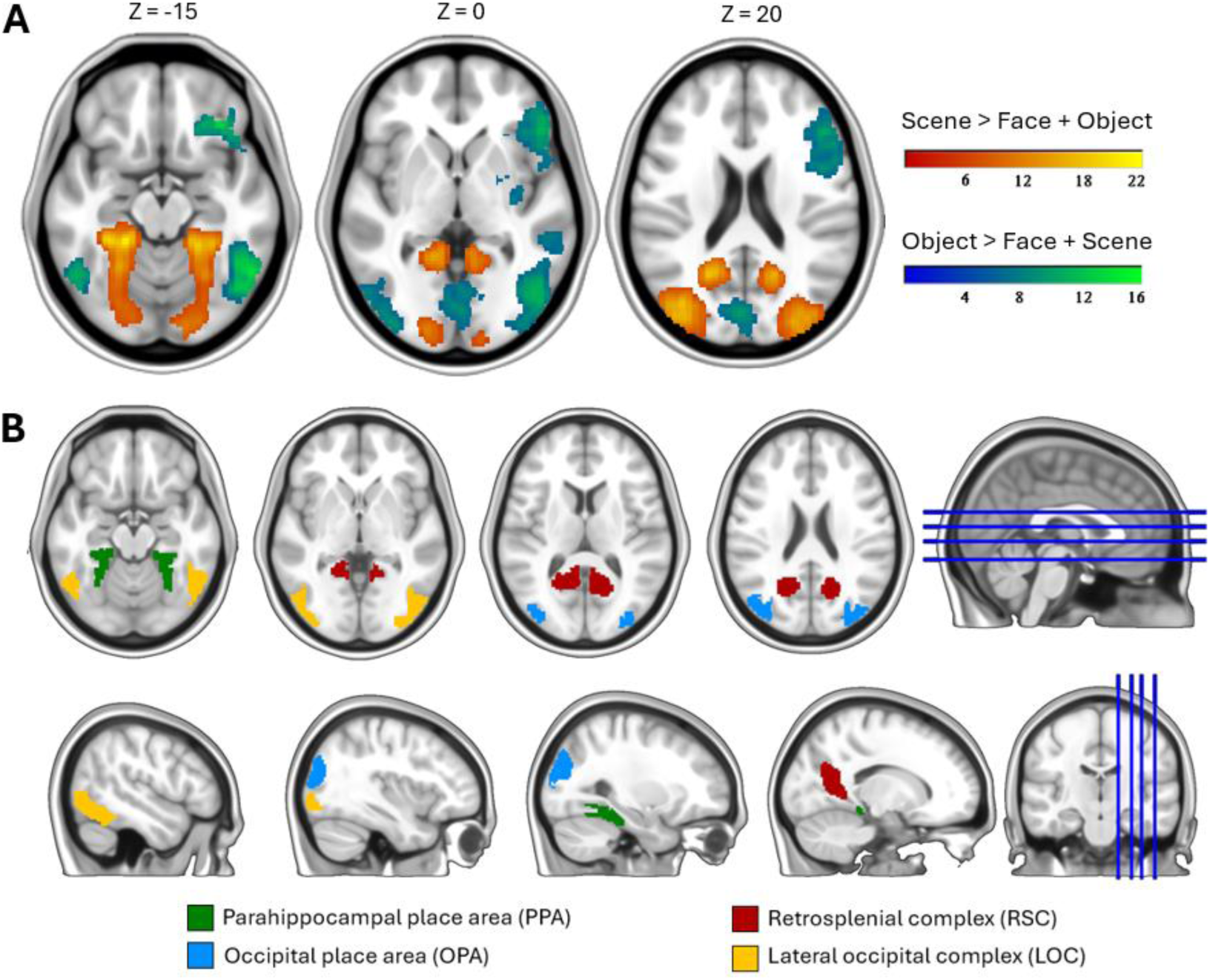
**(A)** Scene-selective and object-selective effects identified in the functional localizer task. **(B)** Regions of interest employed in the primary analyses.

To complement hypothesis-driven analyses in these ROIs, exploratory analyses were performed using the Schaefer-1000 whole-brain parcellation to address where in the brain, beyond the four ROIs, can we identify A) age differences in adaptation, B) relationships with spatial memory. These analyses are also confirmatory in nature: if the effects identified in the whole-brain parcellation spatially overlap with a priori ROIs, the analysis provides important reassurance that the conclusions drawn are not dependent on the ROI definition approach. The atlas was resliced to the EPI grid and masked with a binary group mask. Any parcels containing fewer than 10 voxels following reslicing and masking were eliminated, resulting in a total of 990 parcels. Lastly, we performed an exploratory analysis examining functional connectivity between the hippocampus and the visual ROIs (see *Analysis of hippocampal connectivity* below). The anterior (head) and posterior (body & tail) segments of the hippocampus were extracted from the Allen Human Brain atlas in MNI space.

### Trial-wise fMRI BOLD response estimates

Functional data from the spatial memory task were subjected to a ‘least-squares-separate’ GLM analysis (Mumford et al., 2014), to estimate the BOLD response elicited by each image stimulus. A given stimulus image was modeled by a separate GLM using a 5 s boxcar regressor tracking the image presentation and convolved with a canonical hemodynamic response function. The remaining trials were collapsed into two nuisance regressors corresponding to first- and second-presentation events, excluding the specific event being modeled. Six motion regressors reflecting translational and rotational displacement, along with the mean signal of each session, were included as additional covariates of no interest. For additional motion filtering, we eliminated beta estimates that met criteria for ‘motion spikes.’ For each beta estimate, we examined the framewise displacement (FD) of 6 volumes that coincided with the presentation of the stimulus image (first volume = image onset). To ascertain that the data was not excessively scrubbed for motion in older adults, beta estimates were excluded if any of the 6 volumes exceeded FD of 1.2mm.

In the first iteration of fMRI analyses, condition-dependent beta estimates were extracted from each ROI and averaged across trials where the layout was unchanged. This analysis provided the data necessary to statistically compare whether repeated or rotated scenes exhibited an attenuated response in fMRI BOLD relative to novel scenes. Next, to better control for potential age differences in the gain of the HRF, a metric of adaptation was computed for each subject, using a formula which scales the difference between novel and second presentation betas by the pooled standard deviation: **adaptation = (μβ *_novel_* – μβ *_second_*) / (√(δ *_novel +_* δ *_second_*))/2**. In analyses of raw beta parameters, only trials with the same (unchanged) configuration were included to isolate effects specific to changes in perspective. Importantly, the outcomes of the analyses did not change with the inclusion of all trials. Accordingly, the scaled metric of adaptation includes trials irrespective of layout change to double the statistical power available for trial-wise analyses.

### Time Course Estimation

A finite impulse response (FIR) model was used to visualize the time course of the BOLD response across trials in each of a priori ROI. An FIR model was employed to complement the LSS-based analyses of adaptation by enabling the assessment of condition-specific BOLD dynamics without imposing assumptions about the shape and timing of the hemodynamic response function. Convergent patterns of age effects in univariate LSS and FIR-based analyses would thus suggest that the observed age differences in fMRI BOLD are unlikely to be explained by age differences in HRF timing/shape or differential levels of shared variance between two temporally adjacent event regressors (i.e., first and second presentation of the environment). The time courses were estimated across 14 time points with a sampling interval consistent with TR = 1.52 s. The first sample coincided with the onset of the first stimulus image, and the final sample extended 4 s following the second stimulus image.

### Relationship with spatial memory performance

Generalized linear mixed-effects models (glmer, Bates et al., 2015) were used to examine whether trial-wise estimates of fMRI adaptation relate to the probability of a correct spatial memory response. The models included a trial-wise vector of memory success (correct/incorrect) as the dependent variable. The fixed predictors included the factors of age group, perspective, and trial-wise estimates of adaptation (i.e., mean difference in the response for first vs second stimulus presentation of that trial, scaled by the across-trial pooled standard deviation). The models specified a random intercept on a subject-wise basis, as well as a random slope for adaptation to account for between-subject variability in baseline performance and in the effect of adaptation. Note that the random slope was included in the ROI analysis but not in the analogous whole-brain Schaefer 1000 parcellation analysis to prevent convergence concerns in some of the parcels. In analyses of the four ROIs, the first set of analyses firstly assessed whether adaptation interacted with age group and perspective in predicting spatial memory success. In the presence of an interaction, such models would be followed up with analyses performed separately for the two perspective conditions (in the case of a significant adaptation x perspective interaction), or for the two age groups (in the case of a significant adaptation x age group interaction). In the absence of significant interactions, we conclude that the effect of adaptation on predicting spatial memory success does not differ by perspective or age group, and we evaluate the association between adaptation magnitude and the probability of a correct spatial memory response across all trials while keeping age group and perspective constant.

The analysis above was complemented by an analogous linear mixed effects model, but this time with subject-wise d’ as the dependent variable. This analysis was motivated by the fact that, although the glmer models provide more statistical power by virtue of accounting for variability in spatial performance at the level of individual trials, d’ better accounts for potential age and condition differences in response bias. As previously, age group, perspective, and adaptation were employed as predictors of performance, and relevant adaptation x age group and adaptation x perspective interactions were tested. Any potential follow-up analyses were performed in accordance with the approach described in the glmer methods above.

### Hippocampal beta-series connectivity

Exploratory analyses examining potential age differences in task-evoked functional connectivity between the hippocampus and the four *a priori* ROIs were assessed using a beta-series correlation approach (Rissman et al., 2004). For each participant, trial-wise estimates were obtained using the least-squares-separate approach, as described above. For a given region, we constructed a vector of trial-by-trial mean beta estimates across all betas associated with the second stimulus presentation. Functional connectivity between two regions was operationalized as the Fisher z-transformed Pearson correlation between their respective beta-series. Age differences were evaluated using an independent samples t-test, and linear multiple regression analyses were employed to determine whether hippocampal connectivity relates to fMRI adaptation or estimates of d’, and whether any potential associations differ between age groups.

## Results

### Behavioral analyses reveal age differences in spatial memory and response bias

Hit rates and false alarm rates are summarized in **Table 1**, while d’ and response bias (c) metrics are illustrated in **Figure 3**. D’ values were entered into a 2 (age group) x 2 (perspective) mixed ANOVA, which resulted in a significant age effect (F_(1,49)_ = 23.56, p < 0.001, partial η^2^ = 0.33), a significant effect of perspective condition (F_(1,49)_ = 232.81, p < 0.001, partial η^2^ = 0.83), and a non-significant age x perspective interaction (F_(1,49)_ = 1.15, p = 0.289, partial η^2^ = 0.02). These data indicate that although older adults exhibited overall worse spatial performance, this age effect was equivalent across the two perspective conditions. Therefore, older adults were not disproportionately disadvantaged at recognizing rotated scenes.

**Table 1.**
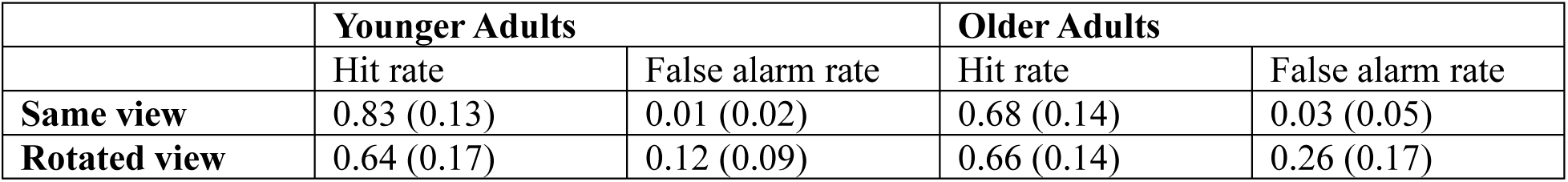
Proportion of hit and false alarm rates calculated by age group and perspective.

|  | Younger Adults |  | Older Adults |  |
| --- | --- | --- | --- | --- |
|  | Hit rate | False alarm rate | Hit rate | False alarm rate |
| Same view | 0.83 (0.13) | 0.01 (0.02) | 0.68 (0.14) | 0.03 (0.05) |
| Rotated view | 0.64 (0.17) | 0.12 (0.09) | 0.66 (0.14) | 0.26 (0.17) |

**Figure 3.**
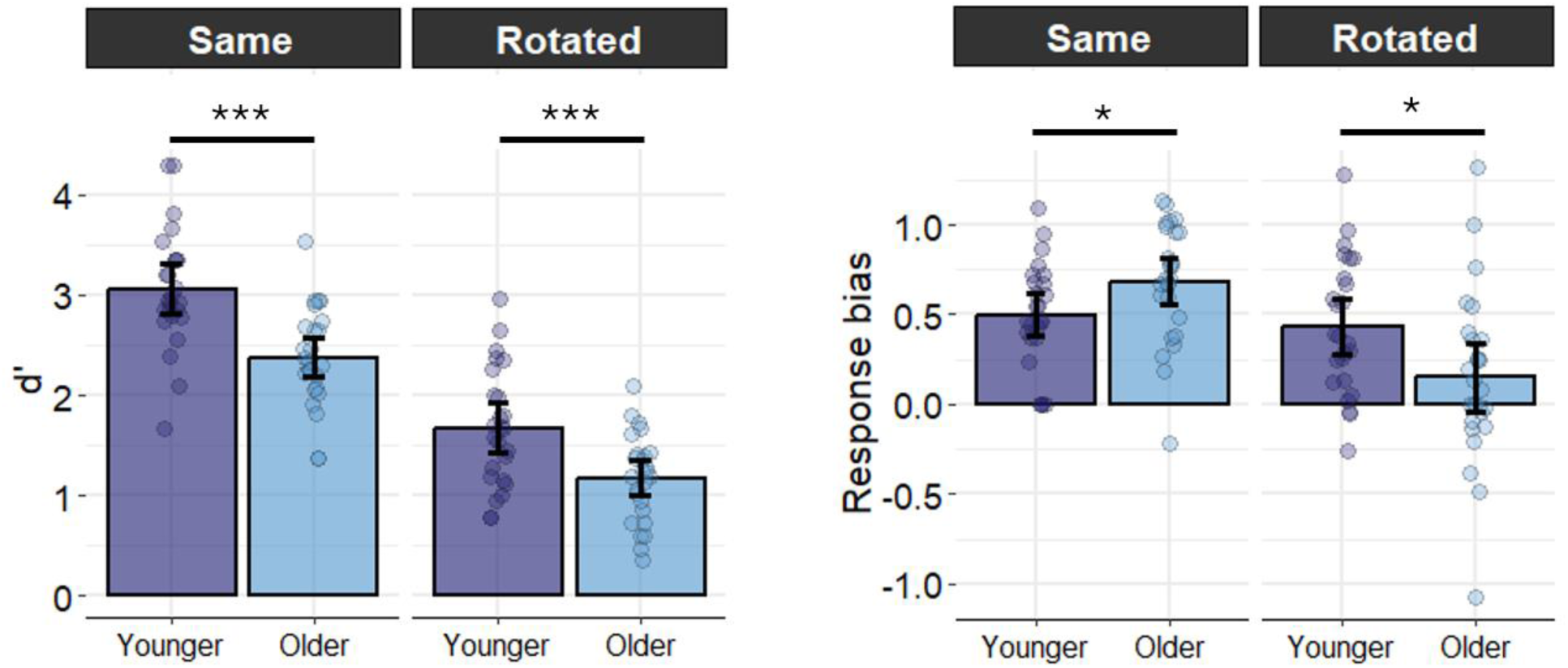
Behavioral performance on the spatial memory task (left = d’; right = response bias). Rotated perspective resulted in worse spatial performance equally in both age groups. However, while younger adults did not exhibit differences in response strategies across perspective conditions, older adults were relatively more conservative for same perspectives and more liberal for rotated perspectives. *** p < 0.001, ** p < 0.01, * p < 0.05; error bars = 95% CI.

Estimates of response bias were entered into an analogous 2 x 2 ANOVA, which revealed a non-significant effect of age group (F_(1,49)_ = 0.28, p = 0.598, partial η^2^ = 0.006), but a significant effect of perspective (F(_1,49)_ = 33.86, p < 0.001, partial η^2^ = 0.41) and a significant age x perspective interaction (F_(1,49)_ = 21.04, p < 0.001, partial η^2^ = 0.30). Follow-up paired t-tests separately in young and older adults revealed that while response bias did not differ by perspective condition in younger adults (t_(24)_ = 0.94, p = 0.359), older adults were significantly more liberal in their responding in the rotated relative to the same perspective condition (t_(25)_ = 6.95, p < 0.001). Additionally, while older adults were more conservative than their younger counterparts during same perspective trials (t_(48.82)_ = 2.15, p = 0.036), they were relatively more liberal than younger adults in their responding during rotated perspective trials (t_(47.30)_ = 2.363, p = 0.022). Taken together, while older adults were not disproportionately impacted by rotated perspectives in the context of d’, they exhibited an age-related shift in decision strategies.

### Older, but not young adults, exhibit adaptation effects to rotated scenes

Parameter estimates from the four *a priori* ROIs were extracted for first presentations (novel images), exact repeats, and rotated repeats. These estimates were then entered into a 2 (age group) x 3 (condition: novel, exact, rotated) x 4 (ROI) mixed effects ANOVA, outcomes of which are reported in **Table 2**. Among the main effects, the ANOVA revealed a null effect of age group, a significant effect of condition, and a significant ROI effect. The ANOVA also resulted in a significant interaction between age group and condition, a condition x ROI interaction, while the age group x ROI and 3-way interactions were not significant. Given that the condition x ROI interaction did not involve the factor of age group, we focus our follow-up analyses on the age group x condition interaction, illustrated in **Figure 4**. The planned tests, as per predictions outlined in **Figure 2C**, were aimed at detecting the extent to which fMRI BOLD exhibits adaptation effect in response to exact and rotated repeats relative to novel images. Therefore, the pairwise tests involved novel vs. exact and novel vs. rotated contrasts across ROIs, which were conducted separately for young and older adults.

**Figure 4.**
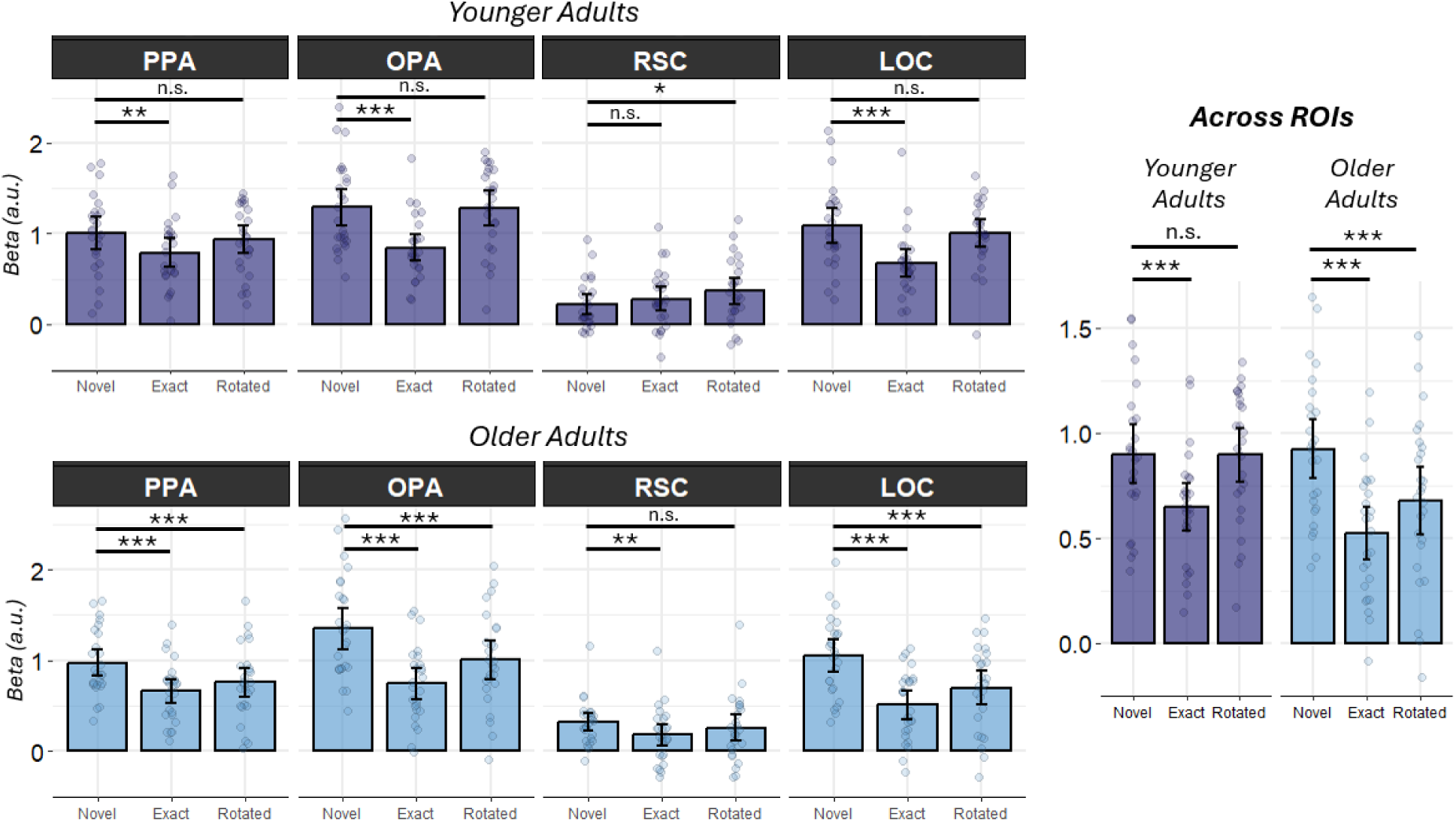
Raw parameter estimates plotted separately for the regressors associated with novel, exact repeats (same perspective) and rotated repeats (rotated perspective). Young adults exhibit adaptation effects, operationalized as the difference between novel – repeat, for exact repeats only. In contrast, older adults exhibit significant adaptation for trials of the same as well as the rotated perspective. *** p < 0.001, ** p < 0.01, * p < 0.05; error bars = 95%CI

In young adults, adaptation effects were evident for exact repeats (t_(24)_ = 3.90, p = 0.001), but not for rotated repeats (t_(24)_ = 0.08, p = 0.939). In contrast, both exact and rotated repeats elicited an attenuated response in older adults (t_(25)_ = 8.76, p < 0.001, and t_(25)_ = 5.54, p < 0.001, respectively). Notably, while there were no age differences in the fMRI response elicited by novel images (t_(48.97)_ = 0.243, p = 0.809), nor exact repeats (t_(48.77)_ = 1.544, p = 0.129), responses elicited by rotated images were significantly lower in older than younger adults (t_(47.56)_ = 2.219, p = 0.033), consistent with the notion of an age-related increase in adaptation effects for rotated scenes. The results of these analyses align with our pre-experimental predictions outlined in **Figure 2C** whereby adaptation effects for exact repeats are evident in both age groups, but only in the older age group for rotated repeats.

### Differential patterns of adaptation effects reflect reduced fMRI responses during second presentation in older adults, regardless of condition

Building on the observations above, we next compared repetition effects across conditions using an index of adaptation magnitude (see Methods). This measure facilitates a direct comparison of adaptation magnitude between exact and rotated stimuli, while explicitly accounting for the response elicited by novel stimuli. Adaptation metrics were entered into a 2 (age group) x 2 (condition) x 4 (ROI) ANOVA (**Table 2**), resulting in a significant effect of age group, condition, and ROI. The age x ROI and condition x ROI interactions were significant, but critically, the ANOVA resulted in non-significant age x condition and 3-way interactions. As illustrated in **Figure 5A**, although the data reported above indicate that older adults, but not younger adults, exhibit repetition effects for both exact and rotated trials, this pattern appears to be largely explained by an overall reduction in the BOLD response magnitude across conditions, rather than a condition-specific age difference in adaptation.

**Table 2.** Statistical outcomes of Age group x Condition x ROI ANOVAs on raw parameter estimates and the scaled adaptation metric.

| *Parameter estimates* |  |  |  |  |
| --- | --- | --- | --- | --- |
| | df | F-value | partial $\eta^2$ | p-value |
| Age group | 1, 49 | 1.60 | 0.032 | 0.212 |
| Condition | 1.7, 84.1 | 47.28 | 0.491 | < 0.001 |
| ROI | 2.4, 116.5 | 110.95 | 0.694 | < 0.001 |
| Age x Condition | 1.7, 84.1 | 6.42 | 0.116 | 0.004 |
| Age x ROI | 2.4, 116.5 | 0.66 | 0.013 | 0.545 |
| Condition x ROI | 3.7, 178.7 | 44.04 | 0.473 | < 0.001 |
| Age x Condition x ROI | 3.7, 178.7 | 2.12 | 0.041 | 0.086 |
| *Adaptation magnitude* |  |  |  |  |
| | df | F-value | partial $\eta^2$ | p-value |
| Age group | 1, 49 | 23.78 | 0.327 | < 0.001 |
| Condition | 1, 49 | 33.35 | 0.405 | < 0.001 |
| ROI | 2.4, 118.5 | 39.32 | 0.445 | < 0.001 |
| Age x Condition | 2.4, 118.5 | 0.02 | 0.001 | 0.889 |
| Age x ROI | 2.4, 118.5 | 9.41 | 0.161 | < 0.001 |
| Condition x ROI | 2.4, 118.5 | 25.28 | 0.340 | < 0.001 |
| Age x Condition x ROI | 2.4, 118.5 | 2.64 | 0.051 | 0.065 |

Time-course data from the FIR models are illustrated in **Figure 5B**. Both age groups exhibited a peak in the FIR time-course approximately at the 5^th^ timepoint for the first stimulus presentation, and at the 13^th^ timepoint for the second stimulus presentation (i.e., in both cases, 6 seconds post-stimulus onset). Therefore, we examined the effects of perspective and age group on the average peak response across timepoints 4-6 for the first presentation, and 12-14 for the second presentation. Across ROIs, no effects of age group, perspective, nor their interaction were evident for timepoints 4-6 (age group: F_(1,49)_ = 0.99, p = 0.324, partial η^2^ = 0.02; perspective: F_(1,49)_ = 0.44, p = 0.509, partial η^2^ = 0.01; interaction: F_(1,49)_ = 2.97, p = 0.091, partial η^2^ = 0.06). For timepoints 12-14, the age group effect was significant (F_(1,49)_ = 4.38, p = 0.042, partial η^2^ = 0.08), as well as the effect of perspective (F_(1,49)_ = 22.90, p < 0.001, partial η^2^ = 0.318). The perspective effects, however, were not moderated by age group (F_(1,49)_ = 0.02, p = 0.897, partial η^2^ = 0.001). Taken together, the FIR analyses confirm that older adults exhibited an attenuated fMRI BOLD response during the presentation of the second stimulus *regardless of condition*, and that this attenuation is statistically equivalent for both same and rotated perspectives. Importantly, given the absence of age differences during the first stimulus presentation, this age effect is unlikely to be caused by a global stimulus-independent reduction in the hemodynamic response.

**Figure 5.**
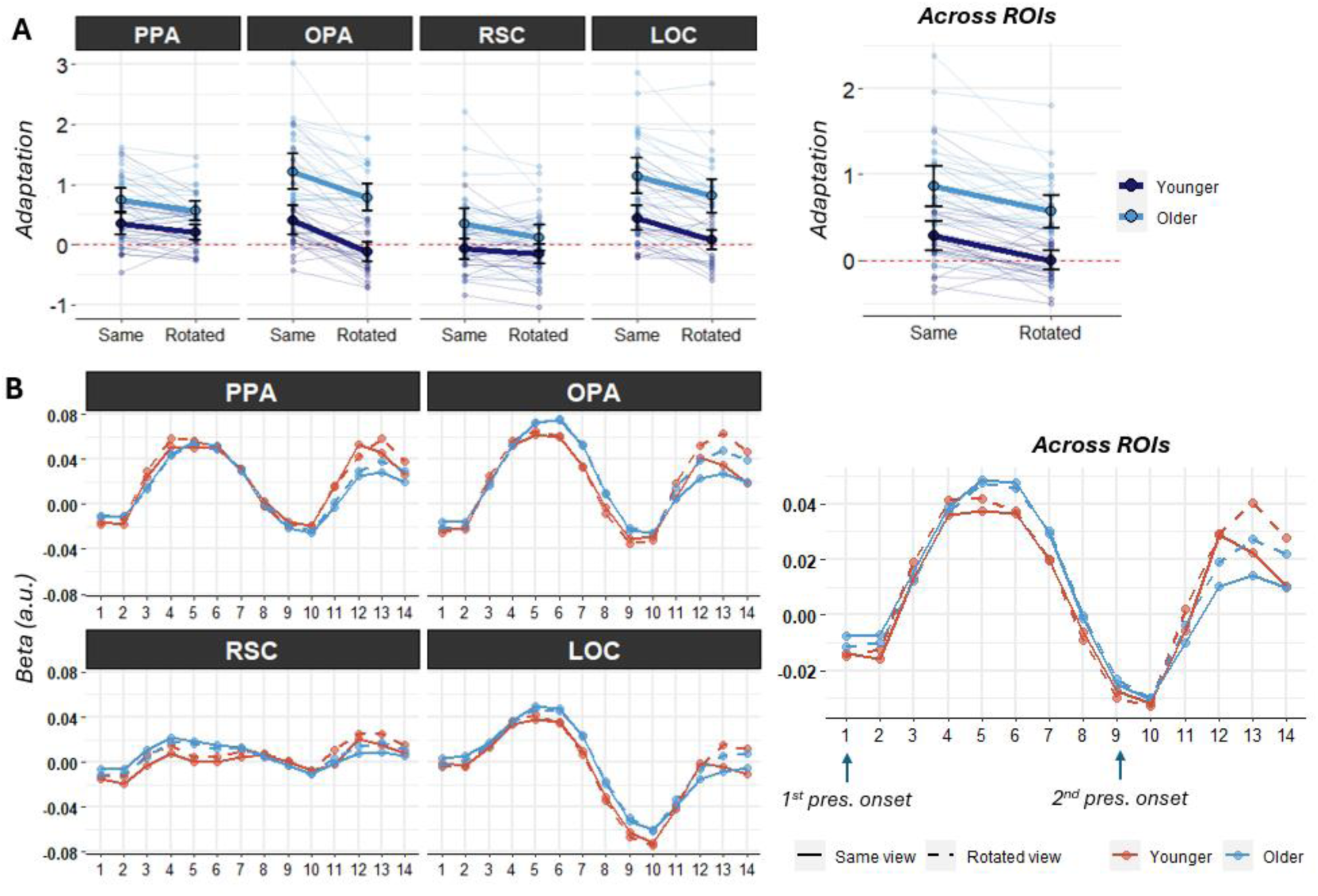
**(A)** fMRI adaptation metric summarized by age group and perspective, plotted separately for the four ROIs (left) as well as averaging across the ROIs (right). The data reveal that while adaptation differs between age groups and perspective conditions, there is a clear absence of an interaction between age group and perspective. Error bars = 95% CI **(B)** Time-course data obtained from FIR GLMs. The data confirm that the fMRI BOLD response is modulated by perspective equally in both age groups, but older adults demonstrate overall greater adaptation (lower fMRI BOLD).

### Relationship between adaptation and spatial performance

In the next series of analyses, generalized linear mixed-effects models (*glmer*) were used to examine whether trial-wise estimates of fMRI adaptation relate to the probability of a correct spatial memory response. In neither of the four ROIs did the factor of adaptation interact with age group (ps > 0.140), indicating that any potential associations with spatial memory success were age-invariant. Perspective also did not interact with adaptation in any of the ROIs (p > 0.318; except for a trend in the OPA at p = 0.052, where adaptation was a weaker but still significant predictor of a correct response for rotated (z = 2.771, p = 0.005) relative to same perspective trials (z = 4.329, p < 0.001). Final models including age group, fMRI adaptation, and perspective revealed that, in all cases, age group and perspective were highly predictive of memory success, consistent with prior behavioral analyses (ps < 0.001). With age group and perspective held constant, adaptation was a significant predictor in PPA (z = 2.425, p = 0.015), OPA (z = 5.051, p < 0.001), and LOC (z = 3.744, p < 0.001), while the association was only trending in the RSC (z = 1.800, p = 0.072). Resulting odds ratios are illustrated in **Figure 6A** and **Table 3**. These analyses suggest that greater adaptation positively covaries with the probability of a correct spatial memory response across perspective conditions and across age groups.

**Table 3.** Odds ratios (95% CI) resulting from the generalized linear mixed-effects models.

|  | PPA | OPA | RSC | LOC |
| --- | --- | --- | --- | --- |
| **Perspective** | 2.58 [2.24, 2.97] | 2.47 [2.15, 2.85] | 2.59 [2.25, 2.98] | 2.54 [2.20, 2.92] |
| **Age group** | 1.64 [1.33, 2.03] | 1.77 [1.42, 2.20] | 1.61 [1.29, 2.00] | 1.71 [1.38, 2.12] |
| **Adaptation** | 1.07 [1.01, 1.13] | 1.14 [1.08, 1.19] | 1.05 [1.00, 1.11] | 1.11 [1.05, 1.17] |

**Figure 6.**
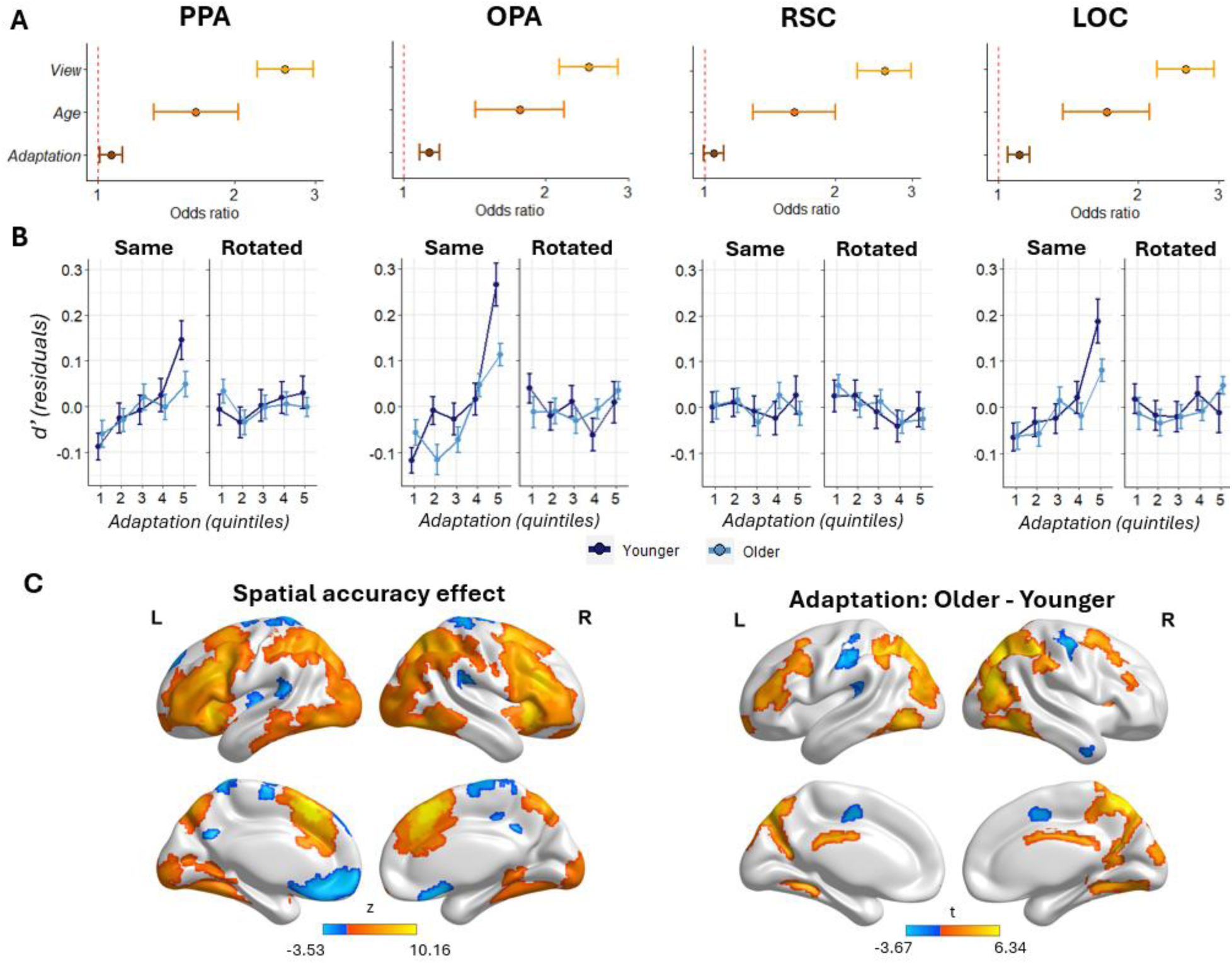
**(A)** Odds ratios and Wald confidence intervals from the generalized linear mixed-effects models were obtained by exponentiating fixed effects coefficients. Odds ratios greater than 1 indicate that the odds of a correct spatial response are higher for the test condition relative to the reference level. References in the binary factors were set to the rotated perspective (OR > 1 equals higher likelihood for same perspective) and the older adult group (OR > 1 equals higher likelihood younger adults). Across PPA, OPA, and LOC, higher adaptation covaries with higher probability of a correct spatial memory judgement while holding perspective and age group constant. **(B)** Increasing adaptation (illustrated as quintiles) is predictive of higher d’ across both age groups, but only for trials of the same perspective. Here, d’ is plotted in the form of residuals after regressing out variance associated with age group. **(C)** Whole-brain results illustrating regions where adaptation was associated with probability of a correct spatial memory judgement (left; orange = higher adaptation predicts higher probability of a correct judgement), and regions where age differences in adaptation were present (right; orange = higher adaptation [lower fMRI BOLD] in older adults).

We also aimed to confirm these conclusions outside of our four *a priori* ROIs with a whole-brain exploratory analysis, where an analogous glmer model was applied to each of the Schaefer 1000 cortical parcels. The outcomes, illustrated in **Figure 6C** (significant at FDR-corrected p < 0.05), reveal positive associations (greater adaptation = higher likelihood of a correct response) across the following regions: early visual cortex, inferior and lateral occipito-temporal cortex, superior occipital cortex, supramarginal and inferior parietal gyrus, middle and inferior frontal gyrus, anterior insula. Negative associations were present in a few parcels in the superior parietal cortex, precuneus, and medial prefrontal cortex. For completeness, **Figure 6C** also illustrates the regions for which age differences in adaptation were present in a linear mixed-effects model when variance associated with perspective was accounted for (*adaptation ∼ age + perspective;* FDR-corrected p < 0.05). As is evident, many of the same regions which covaried with spatial memory success were also associated with significant age differences in adaptation.

Although the probability of a correct response provides more power to estimate associations between trial-level measures of adaptation and accuracy, it differs from overall discriminability (d’) in that it is strongly dependent on bias. This is an important consideration as behavioral analyses reported above revealed that bias differed by age group and perspective condition. For that reason, we complemented the models reported above with linear mixed-effects analyses which employed d’, age group, and perspective as predictors of adaptation. The association between d’ and adaptation differed by perspective in all four ROIs (ps < 0.01), but not by age group (ps > 0.290). Therefore, separate analyses were performed for trials of the same and rotated perspectives using d’ and age group as predictors. For the same perspective trials, d’ covaried positively with adaptation in the PPA (t_(49.24)_ = 2.188, p = 0.033), OPA (t_(48.60)_ = 3.384, p = 0.002), LOC (t_(48.68)_ = 2.279, p = 0.027), but not in the RSC (t_(49.16)_ = −0.116, p = 0.909). No significant associations were evident for rotated perspectives (ps > 0.245). To summarize, greater adaptation shows an age-invariant association with better spatial memory performance, but only in the same perspective trials when response bias is accounted for.

### The strength of hippocampal connectivity with visually responsive cortical ROIs does not differ by age but is predictive of spatial memory accuracy

Given the hippocampus’ role in spatial memory and its sensitivity to increasing age (see Introduction), we performed a series of exploratory analyses examining whether age differences in adaptation magnitude relate to potential age differences in hippocampal connectivity with the four ROIs. No age differences in connectivity were evident between either the anterior (head) or posterior (body + tail) segments of the hippocampus and the ROIs (ps > 0.2, **Figure 7**). Given that connectivity metrics were highly correlated across subjects, and to reduce the number of multiple comparisons, we averaged across ROIs to obtain average hippocampal connectivity within visually responsive cortical regions, and we tested whether it covaried with adaptation or estimates of d’ using two multiple regression models. In neither case did we find a significant interaction with age group (adaptation: t_(47)_ = 0.101, p = 0.920; d’: t_(47)_ = 0.377, p = 0.738), indicating that any potential relationship was, once again, age-invariant. A partial correlation between hippocampal connectivity and mean adaptation, while covarying out the factor of age group, revealed a negative association between the two variables (r_partial_ = −0.334, p = 0.018). A similar negative association was also identified in an analogous partial correlation analysis between hippocampal connectivity and d’ (r_partial_ = −0.475, p < 0.001). The results thus reveal that hippocampal decoupling (lower connectivity) with visually responsive cortical regions during spatial retrieval appears to be beneficial to spatial memory performance. Importantly, this relationship and the magnitude of decoupling did not differ between young and older adults.

**Figure 7.**
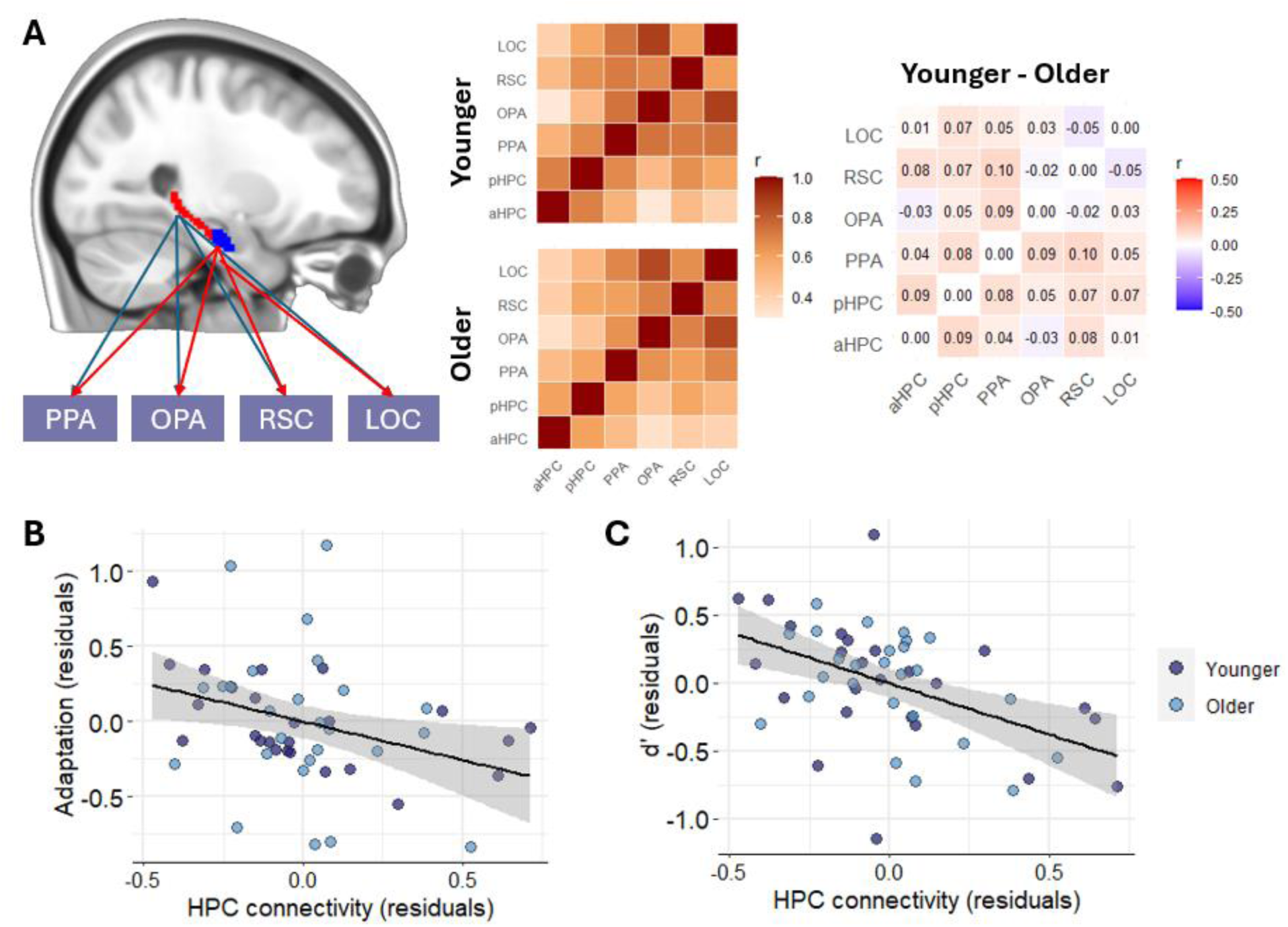
**(A)** Beta-series correlation analysis of functional connectivity was performed across all second image beta estimates, with a particular emphasis on the connectivity between the hippocampus and the four a priori ROIs. The strength of connectivity was equivalent between younger and older adults. For illustration purposes, Fisher-z values were transformed back to Pearson’s r space, and values on the right panel reflect mean differences in Pearson’s r between younger and older adults. None of the cells in the matrix showed significant age differences. **(B)** Mean connectivity between the hippocampus and the four higher-level visual ROIs is negatively correlated with the magnitude of adaptation across subjects. The relationship is age-invariant, and the values are plotted after residualizing variance associated with age group. **(C)** Mean hippocampal connectivity is also negatively associated with d’. This relationship is also age-invariant.

## Discussion

The present study examined age differences in viewpoint specificity as a potential neural mechanism contributing age-related decline in spatial memory. To summarize, behavioral analyses revealed lower spatial memory performance in older adults across both the same and rotated perspective conditions. Raw fMRI parameter estimates revealed an attenuated fMRI response (adaptation) to the same but not rotated perspectives in young adults, but to both perspective types in older adults, consistent with a potential altered adaptation response to rotated scenes with age. When adaptation was quantified as a scaled difference metric, however, older adults showed a general attenuation of the fMRI response (higher adaptation) regardless of the perspective condition. Interestingly, although older adults showed higher adaptation and worse spatial accuracy, higher trial-wise estimates of adaptation were associated with a higher likelihood of correct spatial memory independent of age. These findings highlight that adaptation reflects at least two independent sources of variance: spatial memory across the lifespan, as well as variance that is independently attributable to increasing age. Below, we explore the significance of these findings for the understanding of the mechanisms that underpin age differences in spatial memory and navigation.

Turning first to behavioral findings, we found that older adults performed worse on the spatial memory task compared to younger adults across both perspective conditions. Although the absence of an age x condition interaction is inconsistent with the notion that aging disproportionately impacts spatial memory when invoking viewpoint-invariant (allocentric) representations (e.g., Bohbot et al., 2012; Lithfous et al., 2014; Castegnaro et al., 2025), our findings are consistent with behavioral studies that employ similar tasks (Montefinese et al., 2015; Muffato et al., 2019; Hilton et al, 2020; Segen et al., 2021; but see Segen et al., 2022). That said, while young adults did not change their decision strategies (response criterion) by condition, older adults were more conservative than their younger counterparts for the same perspectives and relatively more liberal for rotated perspectives. This indicates that lower spatial accuracy in older adults may be attributable to at least two distinct cognitive processes. For the same perspective, one possibility is that a more conservative response threshold may reflect reduced confidence in the context of relatively higher visual similarity between first and second presentations of the environment, pointing to weaker (or less precise) mnemonic evidence available for perceptual matching. As for the rotated perspectives, the liberal criterion appears to be primarily reflective of an elevated false alarm rate (endorsing an unchanged layout ‘different’). The rotated condition requires mentally transforming the viewpoint and matching it to the encoded stimulus. If this transformation is noisy, the internally rotated representation will not perfectly match the initially experienced viewpoint. Taken together, age differences in the present spatial memory task may reflect noisy relational comparison (less precise mnemonic evidence and noisier viewpoint transformation) as opposed to a selective deficit in allocentric strategies.

Viewpoint specificity was examined by evaluating the magnitude of fMRI adaptation to the same versus rotated perspectives. The younger adult cohort exhibited adaptation to same but not rotated perspectives, aligning with past studies suggesting viewpoint specific codes in areas like parahippocampal gyrus (e.g., Epstein et al., 2003, 2007, 2008). These data are interpreted as evidence that, across the four a priori ROIs, the environments are represented in the context of their perceptual appearance, as opposed to a holistic abstract map of the spatial layout. For older adults, one possible interpretation for the finding that adaptation effects were observed for both the same and rotated viewpoints is that their representations were *less* viewpoint specific. Such an interpretation would point to an age-related reduction in neural specificity of scene processing, possibly due to greater noisier underlying population activity. That said, an alternative explanation pertains to an age-related shift towards greater viewpoint tolerance (rather than reduced viewpoint-specificity), which assumes that scene representations during retrieval may increasingly abstract away high-fidelity perceptual detail in favor of viewpoint generalization (e.g., Steel et al., 2021; Bainbridge et al., 2021; Srokova et al., 2022). Given that real-world environments are rarely encountered from a single vantage point, such generalization may be compensatory by allowing older individuals to better recognize the same place despite changes in perspective.

Although the pattern observed in the raw parameter estimates are consistent with these interpretations, the results rely on differences in the absolute response magnitude of the BOLD signal and may therefore conflate viewpoint specificity with overall differences in adaptation. For that reason, we also employed a scaled difference metric of adaptation which explicitly quantifies the difference in fMRI responses between the first and second presentations of the environment. This analysis revealed overall greater adaptation in older than younger adults, but critically, an absence of an age x condition interaction, highlighting that the magnitude of adaptation did not differ between the same and rotated perspectives. This is a critical consideration: although the raw fMRI response suggested greater adaptation for rotated scenes, the scaled analysis indicated that older adults showed greater adaptation across *both* conditions.

There are two different classes of accounts which may elucidate this result: physiological (vascular) and cognitive. First, reduced fMRI response in older age may reflect reduced HRF gain or slower hemodynamic recovery (D’Esposito et al., 1999; Huettel et al., 2001; West et al., 2019). A broader or slower HRF in aging, however, would predict an increase in the shared variance between the regressors in the GLM, which would affect the estimation of both events simultaneously rather than selectively suppressing only one of them. Therefore, given the absence of an age effect for the first presentation of a scene, physiological mechanisms cannot fully account for the observed results. While the present data do not allow us to adjudicate the cognitive source, two possibilities warrant consideration. First, it is likely that, when viewing the environment for the second time, participants reinstate the encoded representation of the environment to perceptually match it to the stimulus they are currently viewing. Considering prior evidence for age-related differences in the mnemonic reinstatement of previously experienced stimuli (for review, see Rugg & Srokova, 2024), the observed effect may reflect an attenuation in retrieval-related activity during the second presentation of the environment. Alternatively, or in addition, the task requires participants to re-process and update the scene representation when they view it again, whereby spatial relationships between objects need to be re-evaluated to make a spatial judgement. Older adults may engage less in such representational updating (Wahlheim & Zacks, 2019, Merhav & Wolbers, 2019, Qin & Basak, 2020), leading to attenuated responses in regions typically recruited during visual processing.

We also found an association between fMRI adaptation and spatial memory performance across both age groups, although this effect was relatively more robust for trials of the same perspective. It is notable that the direction of the brain-behavior association was the opposite of the age effect discussed above: while older adults showed greater adaptation and poorer spatial memory, greater trial-wise estimates of adaptation were predictive of better spatial accuracy across both age groups. As noted previously, it is unlikely that these two effects reflect the same cognitive process. Mirroring the spatial updating account discussed above, one possibility is that the variance sensitive to cognition is associated with more efficient re-processing of the environment when it is viewed for the second time (instead of weaker updating, as may be the case for the age effect). Because minimal spatial updating is necessary when the internal representation of the encoded environment is strong, fewer computational resources are recruited, leading to lower fMRI BOLD.

Lastly, we performed exploratory beta-series correlation analysis between hippocampal and higher-level visual ROIs, motivated by the notion that age differences in spatial memory may stem from differences in cortical activity received by the hippocampus. We found that the strength of trial-by-trial coupling between visual regions and the hippocampus was comparable between young and older adults. Additionally, lower covariance was associated with higher adaptation and better spatial memory across age groups. This suggests that when a stable representation is present in visual cortex, hippocampal engagement may be less critical for successful performance.

An important limitation of the present study lies in its cross-sectional design. It is possible that the present results may be confounded by age differences in prior experience with visual desktop virtual reality settings, such as video games (Charness & Boot, 2022). Additionally, in real world contexts, spatial memory and navigation involves active ambulation, which enables the updating of the spatial representation through proprioceptive and vestibular signals (i.e., self-motion cues; Hejtmanek et al., 2020; Holmes et al., 2018; Richardson et al., 1999; Ruddle & Lessels, 2006). Indeed, in the context of the present task, spatial memory is more accurate if self-motion cues are available to support performance (Srokova et al., 2026), and the presence of self-motion cues may ameliorate age differences in spatial accuracy (Hill et al., 2024, 2025). It is unknown how such multimodal input may alter brain activity in the context of fMRI.

In summary, the present study provides novel evidence elucidating the fMRI correlates underlying age differences in spatial accuracy for same and rotated perspectives. Although raw estimates pointed to a reduced viewpoint specificity in older adults, a standardized adaptation metric indicated an overall increase in repetition-related adaptation across both perspective conditions. The pattern of fMRI results mirrors behavioral findings in which older adults exhibit reduced accuracy across both perspective conditions. Overall, the fMRI and behavioral data argue against an age-related deficit that would be specific to viewpoint-invariant (i.e., allocentric) scene representations, and caution against interpreting greater adaptation in older adults as a marker of diminished viewpoint specificity.

## Acknowledgements

This work was supported by NIH/NIA grant R01AG003376. S.S. is supported by the Postdoctoral training program of the Arizona Alzheimer’s Consortium, NIH/NIA grant T32AG044402.

## Author contributions

S.S. and A.D.E designed research, A.D.E. and C.A.B. acquired funding, S.S. performed research, S.S. analyzed data, S.S. and A.D.E wrote the paper, C.A.B. edited the paper.

